# Generating effective models and parameters for RNA genetic circuits

**DOI:** 10.1101/018358

**Authors:** Chelsea Y. Hu, Jeffrey D. Varner, Julius B. Lucks

## Abstract

RNA genetic circuitry is emerging as a powerful tool to control gene expression. However, little work has been done to create a theoretical foundation for RNA circuit design. A prerequisite to this is a quantitative modeling framework that accurately describes the dynamics of RNA circuits. In this work, we develop an ordinary differential equation model of transcriptional RNA genetic circuitry, using an RNA cascade as a test case. We show that parameter sensitivity analysis can be used to design a set of four simple experiments that can be performed in parallel using rapid cell-free transcription-translation (TX-TL) reactions to determine the thirteen parameters of the model. The resulting model accurately recapitulates the dynamic behavior of the cascade, and can be easily extended to predict the function of new cascade variants that utilize new elements with limited additional characterization experiments. Interestingly, we show that inconsistencies between model predictions and experiments led to the model-guided discovery of a previously unknown maturation step required for RNA regulator function. We also determine circuit parameters in two different batches of TX-TL, and show that batch-to-batch variation can be attributed to differences in parameters that are directly related to the concentrations of core gene expression machinery. We anticipate the RNA circuit models developed here will inform the creation of computer aided genetic circuit design tools that can incorporate the growing number of RNA regulators, and that the parameterization method will find use in determining functional parameters of a broad array of natural and synthetic regulatory systems.

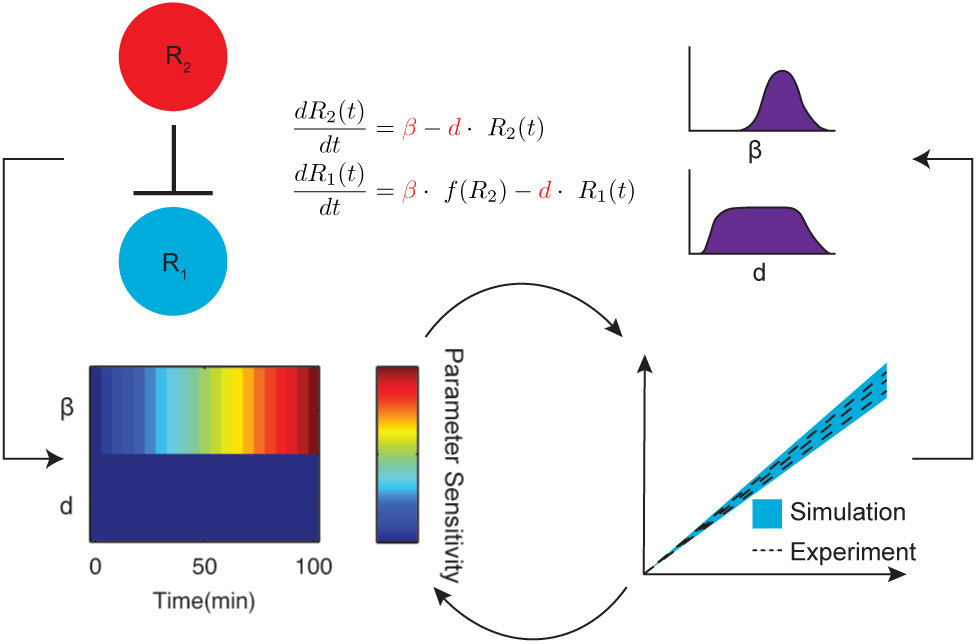

## Introduction

A central goal in synthetic biology is to engineer biological systems to optimally perform natural and sometimes novel tasks. These tasks are varied, and are often related to important challenges in sustainability and health. For example, there has been a great deal of effort placed on harnessing the natural capabilities of cells to synthesize complex molecules from renewable feedstocks, and to sense and respond to dynamically changing environments. These capabilities are themselves implemented through genetic circuits – networks of interacting gene expression regulators that can dynamically balance metabolic pathways^1^ and integrate multiple signals to make behavioral decisions^2^, among many other behaviors related to cell replication and survival^3^. Thus a cell’s behavior is intimately connected to the structure of its genetic circuits^1,4^, making the precise engineering of genetic regulators and networks a central goal of synthetic biology^5^.

Over the past decade, synthetic biology has seen significant advances in the quantity and sophistication of engineered genetic circuits^6^. Recently, regulatory RNAs have emerged as powerful components of the synthetic biology toolbox for constructing genetic circuits that control the timing and amount of gene expression^7^. These natural and synthetic RNAs are diverse, and can regulate transcription, translation and RNA degradation, especially in bacteria^8^. There is also a growing body of work on developing design principles for engineering the functional properties of regulatory RNAs. This work makes use of a powerful suite of computational^9^ and experimental^10^ methods to discern the underlying RNA structures behind the regulatory function^7^, and is leading to new classes of highly-designable RNA regulators^11^.

Of particular interest are RNA mechanisms that regulate transcription. These regulators are special because they can be wired together into RNA-only genetic circuits that propagate information as RNAs without the need to translate or degrade intermediate regulatory proteins^12^. Recently it has been shown that RNA transcriptional repressors, also known as attenuators^13^ can be configured into NOR logic gates and transcriptional cascades^12^. They can also be used to construct more sophisticated circuits, such as single-input modules^14^ that sequentially activate the expression of multiple target genes^15^. Advances in RNA engineering approaches have also greatly expanded the types of circuits that can be built out of RNA mechanisms. For example, the creation of small transcription activating RNAs, or STARs, allows the creation of new types of RNA logic gates that implement AND and NIMPLY^16^. This, combined with exciting new developments in using the RNA-protein hybrid Clustered Regulatory Interspaced Short Palindromic Repeats (CRISPR) interference (CRISPRi) system to construct transcriptional repressors^17^, activators^18^, and NOT and NOR logic gates in cells^19^, has started to draw attention to RNA-based genetic circuits as a platform for precisely controlling gene expression.

A major gap in our RNA regulatory toolbox is the lack of a computational framework that can be used to model and ultimately design RNA genetic circuitry. Such a modeling framework is also necessary for incorporating RNA regulators and circuits into a growing suite of computer-aided design (CAD) tools that allow users to use high-level cellular behavioral specifications to design synthetic genetic circuits and select genetic regulatory parts that implement those behaviors20-22. While there has been progress in modeling the impact of individual RNA regulators on tuning gene expression^23-25^, there has been little work in modeling how networks of transcriptional RNA regulators impact the coordinated expression of multiple genes. In contrast, for protein-based genetic circuits, systems of ordinary differential equations (ODEs) that model the basic processes of gene expression in a genetic network as coupled chemical reactions^26^ are commonly used to computationally study both natural^27^ and synthetic^28^ genetic networks. However, it is generally not known if simple ODE-based frameworks work for modeling RNA transcriptional circuits, and if they do, what parameter values are needed for accurate prediction of circuit behavior (Figure 1A).

**Figure 1.**
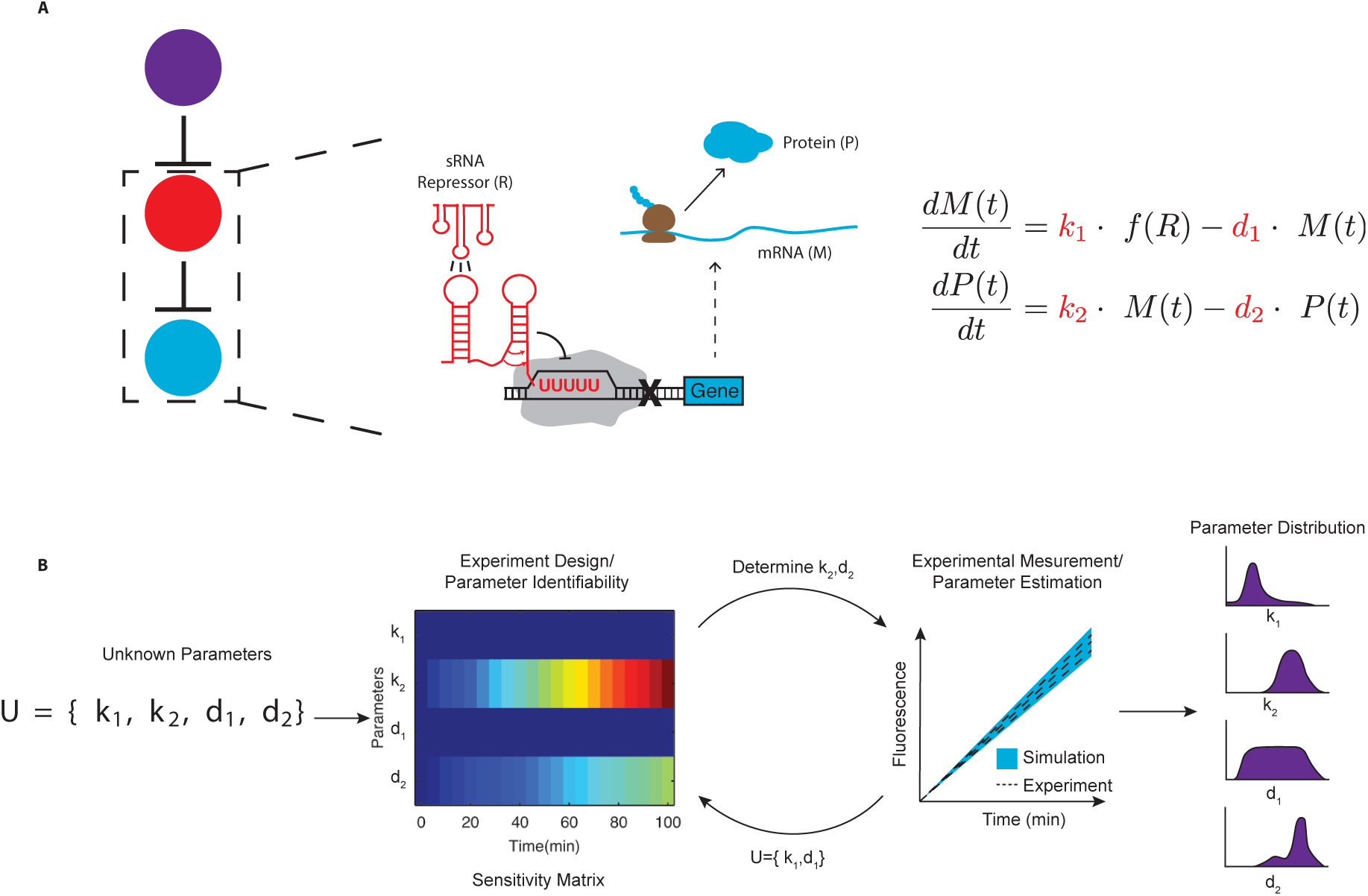
Schematic of the model development and parameterization process. (A) A 3-level, DNA-encoded RNA transcriptional cascade (colored circles) is composed of two orthogonal sRNA repressor/attenuator pairs. The RNAs are configured in a double repression cascade, with the final output being the transcription of a target gene that encodes a translated protein. The complex mechanism of RNA transcriptional repression can be described by coarse-grained ordinary differential equations (ODEs) with a handful of unknown parameters (red). (B) Parameterization experiments can be designed based on a parameter sensitivity analysis of the model equations. This analysis identifies which parameters can be estimated from a particular experimental design. These experiments can then be performed to estimate the indicated parameters. This process can then be iterated until all parameters are estimated, resulting in distributions of parameters that accurately model the desired genetic circuitry.

In this work, we develop an effective modeling framework that quantitatively captures the dynamic outputs of RNA-only transcriptional circuits. To do this, we leverage recent advances in using cell-free transcription-translation (TX-TL) reactions as rapid genetic circuit prototyping environments for characterizing genetic circuit dynamics^15,29,30^ and for modeling gene expression processes^31^. We build this model by studying a double-repression RNA-only transcriptional cascade^12,15^. We start by systematically constructing a system of ODEs that model the expression and degradation of each RNA regulator in the cascade (Figure 1A), and find that RNA maturation delay equations must be added in order to qualitatively capture cascade function. Since many of the model parameters for RNA-only circuits are not available in the literature, we next adopt a systems biology approach to estimate all unknown parameters in this model. Rather than perform a suite of biochemical experiments to measure each parameter in turn, this approach uses sensitivity analysis on the mathematical structure of the genetic network to design a minimal set of characterization experiments that can be used to rapidly and quantitatively determine all parameters in a model^32^ (Figure 1B). Specifically, we show that all thirteen parameters of our model can be estimated from only four TX-TL experiments that can be run in parallel in under two hours. We then use our estimated parameters with the governing ODE framework to predict the function of six new RNA transcriptional circuit variants and show that the simulated predictions compare favorably with experimental measurements. Finally, we perform model parameterization experiments using a different batch of TX-TL reagents, and show that batch-to-batch variation can be attributed to differences in parameters that are directly related to the concentrations of core gene expression machinery. We end with a discussion about how this method can be generalized to rapidly determine parameters for quantitatively modeling the dynamics of synthetic genetic circuits.

## Results and Discussion

### Model Derivation

Our first goal was to develop a modeling framework that could qualitatively capture the dynamical behavior of RNA-only circuits. One mathematical framework that has found wide use in modeling genetic circuits is systems of ordinary differential equations that treat the basic processes of gene expression in a genetic circuit as coupled chemical reactions^27^. While these models can vary in detail, in essence they consider the concentration of a given molecular species in time to be a function of its synthesis and degradation rates^27^ (Figure 1A). These rates in turn can be functions of the concentrations of other regulators in the circuit, effectively coupling the equations according to the circuit’s network topology.

Our goal was to construct the simplest possible model (i.e. fewest number of equations and parameters) that could capture the function of RNA transcriptional repressors and cascades. To do this, we focused on modeling a two-repressor transcriptional cascade that had been previously characterized^12,15^. This cascade was constructed from two orthogonal RNA repressors called transcriptional attenuators (Figure 2A). Transcriptional attenuators act as genetic switches by blocking or allowing transcription through conditional formation of an RNA intrinsic terminator hairpin^33^. In the absence of an antisense sRNA, sequences in the attenuator prevent terminator formation, which allow transcription of a downstream gene. When antisense sRNAs are present, they bind to their attenuator targets, which allows terminator formation and switches transcription to an OFF state.

**Figure 2.**
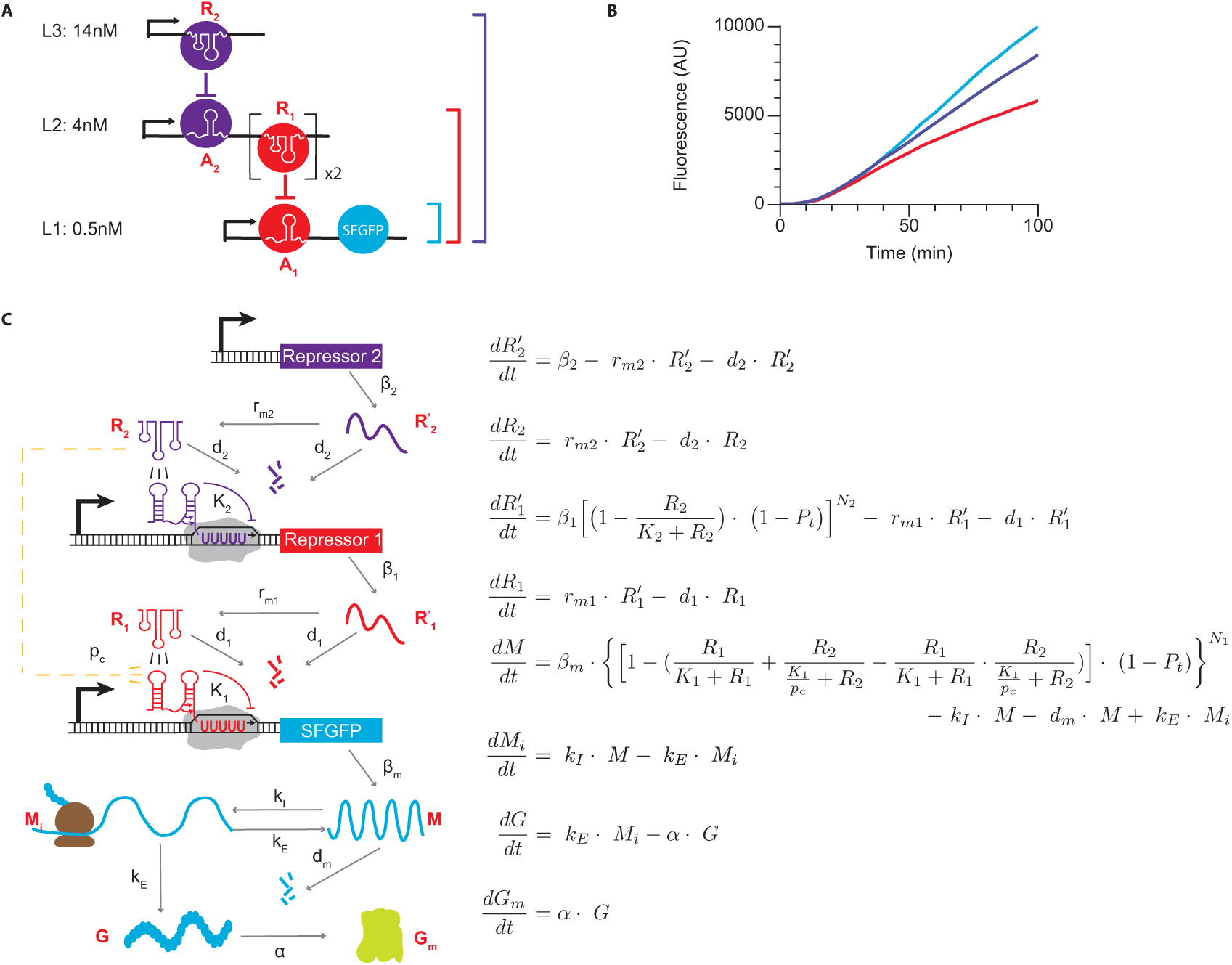
Governing Equations for a 3-level sRNA transcriptional cascade. (A) Schematic of the cascade. Level 1 (L1) plasmid expresses attenuator 1 (A_1_) controlling SFGFP transcription; Level 2 (L2) expresses attenuator 2 (A_2_) controlling the transcription of tandem copies of sRNA repressor 1 (R_1_); Level 3 (L3) expresses sRNA repressor 2 (R_2_). Concentrations of DNA templates used in TX-TL reactions for part B are listed beside the levels. (B) Representative fluorescence signal time trajectories in TX-TL reactions containing three different combinations of DNA templates from the transcriptional cascade in part A. L1 alone leads to a high rate of SFGFP production (blue line); L1+L2 results in results in a reduced SFGFP production rate (red line) due to R_1_ repressing A_1_; L1+L2+L3 (purple line) results in a higher SFGFP production rate than just L1+L2 due to the double negative inversion of the full cascade. (C) Schematic (left) of the mechanistic steps of the cascade in part A that are captured by the governing equations (right). The equations model the tandem copies of R_1_ on L2 as one repressor.

The double-repression RNA transcriptional cascade consisted of three levels^12,15^ (Figure 2A). The bottom level of the cascade (L1) consisted of a constitutive promoter followed by attenuator-1 (A_1_), which controlled the transcription of a downstream super folder GFP (SFGFP)^34^ coding sequence. A_1_ was itself switched to an OFF state by interactions with repressor sRNA 1 (R_1_), which was present in two tandem copies on the second level (L2) of the cascade^12,15^. The complete L2 also contained a constitutive promoter followed by attenuator-2 (A_2_), both upstream of the R_1_ copies. Following previous work, self-cleaving ribozymes were included before each R_1_ copy to ensure proper function^12^. This configuration allowed the transcription of R_1_ to be controlled by repressor sRNA 2 (R_2_), which was expressed from a constitutive promoter on level 3 (L3) of the cascade.

Characterization of the cascade in TX-TL reactions revealed the patterns of fluorescence expected from combining different levels of the cascade in the reactions (Figure 2B). TX-TL reactions consist of three components: cell extract, energy solution/buffer and input circuit DNA. To characterize the performance of RNA genetic circuits, plasmid DNA encoding different combinations of cascade levels were mixed with extract and buffer following previously published protocols^35^ (see Methods). These were then monitored on a plate reader to measure SFGFP fluorescence over time to characterize overall circuit expression. As expected, when only L1 DNA was present, we observed a rapid increase in SFGFP fluorescence, which was decreased when both L1 and L2 DNA were present (Figure 2B). The addition of L3 DNA in the reaction showed an increase over the L1+L2 condition indicating that the double repression of the cascade was functioning properly as had been observed previously^15^.

As a starting point for our model, we considered the expression of each RNA in the system to be governed by one mass balance ODE that accounts for its synthesis and degradation rates. For simplicity, we considered these rates to be described by single constants. To model transcriptional repression, we introduced Hill functions of order one, based on previous experimental work in characterizing sRNA/attenuator transfer functions^12^. This allowed us to construct a set of four equations with eight parameters that captured the flow of information in the RNA transcriptional cascade (SI Equations 1.1-1.4, Figure S1). Note that we incorporated the tandem copies of R_1_ present in L2 of the cascade through the parameters of the model rather than extra mechanistic steps.

Using this simple model, we simulated time dependent trajectories of the cascade (Figure S2). These results showed that there were several qualitative disagreements between the model and the previously reported experimental data for this cascade. In particular, several mechanistic details of RNA transcriptional attenuators were not included in these equations. The most important of these is cross-talk, or the ability for non-designed interactions to cause repression between different levels of the cascade due to the imperfect orthogonality of the repressors used in the system^12^. We incorporated cross-talk into the model by using a constrained fussy logic formulation, treating the bottom level of the cascade as an OR gate module with two signaling species accounting for the contributions of cognate and cross-talk interactions^36^(Figure S1B,SI Equations 2.1-2.4).

Another feature of the RNA transcriptional attenuators not captured is their ability to be placed in tandem next to each other so that the combined attenuator can respond to multiple antisense sRNAs. Previous work had found that attenuators in tandem multiplied their effects and increased their sensitivity to antisense RNA^12^. In fact this feature was exploited in order to create a time-delay in a singe-input module that could sequentially activate the expression of two different genes^15^. To incorporate tandem attenuators in our model, we raised the repressive Hill functions to the power of tandem attenuator number on that level (Figure S1C, S2B, SI Equations 3.1-3.4). Previous work also showed that some aspect of the attenuation mechanism causes repression of the downstream gene even in the absence of any antisense sRNA^12^, which was also incorporated into our model (Figure S1D, S2C, Equations SI 4.1-4.4).

We next began comparing our model to measured fluorescence trajectories from TX-TL reactions (Figure S3). Since we used a fluorescent reporter protein as a final output of the cascade, we added additional equations to model the translation and maturation of the fluorescent reporter protein (SI Equations 5.1-5.6). Specifically, we modeled translation as a 2- step process consisting of initiation and elongation, and ignored SFGFP degradation which is appropriate for TX-TL reactions^35^. During this model formulation process, we noticed that there was a delay between when DNA was introduced in the TX-TL reactions, and the time it took to observe fluorescence that was longer than the expected delay due to SFGFP maturation. Specifically, we were not able to observe a fluorescence signal until ∼20 minutes after the reaction was initiated. As we tried to qualitatively fit the model to the experimental trajectories, we found that this delay was too large to be described by any known mechanism of this system (Figure S3A, B). Thus we hypothesized that the observed delay was specific to the TX-TL system, which could be due to the time needed for activation of extract core machinery after mixing with buffer. To test this, we pre-incubated extract and buffer together at 37°C for 20 minutes before adding DNA and performing the circuit characterization. We found that when extract and buffer were pre-incubated, this delay was removed, allowing greater qualitative agreement with our model (Figure S3C, D).

While we were able to obtain a qualitative match between our model and experimental trajectories, we noticed that when L1 and L2 were both present in TX-TL reactions, we observed a decrease in SFGFP production rate after 30 minutes. This manifested as a downward ‘bending’ of the L1+L2 fluorescence trajectories that we could not qualitatively capture (Figure S4). We hypothesized this was due to the fact that the sRNA R_1_ repressor encoded in the L2 plasmid did not function immediately once synthesized, which was an underlying assumption of the model up to this point. To incorporate this into our model, we added additional sRNA maturation steps into the governing equations (SI Equations 6.1-6.8, Figure 2C). As shown in Figure S2B, after we introduced sRNA maturation delay terms into the governing equations, we were able to qualitatively capture the bending of the L1+L2 trajectories.

Since the same promoter was used on all constructs, β_1_ and β_2_ were determined by multiplying β_m_ by an appropriate factor based on DNA template concentrations. The final governing equation set thus consisted of eight ODEs and thirteen unknown parameters (Figure 2C,Tables 1 and 2), which qualitatively captured the behavior of the three-level RNA transcriptional cascade.

**Table 1.** Model Parameters.

| Model Parameters | Number | Definition | Unit |
| --- | --- | --- | --- |
| $\beta_2$ | 1 | Maximal transcriptional rate of level of $R_2$ (determined from $\beta_m$) | Conc./sec |
| $\beta_1$ | 2 | Maximal transcriptional rate of level of $R_1$ (determined from $\beta_m$) | Conc./sec |
| $\beta_m$ | 3 | Maximal transcriptional rate of M | Conc./sec |
| $K_2$ | 4 | Repression coefficient of $R_2$ on the production of $R_1$ | Conc. |
| $K_1$ | 5 | Repression coefficient of $R_1$ on the production of M | Conc. |
| $k_E$ | 6 | Translational elongation rate of SFGFP | 1/sec |
| $d_2$ | 7 | Degradation rate of the repressor $R_2$ | 1/sec |
| $d_1$ | 8 | Degradation rate of the repressor $R_1$ | 1/sec |
| $d_m$ | 9 | Degradation rate of M | 1/sec |
| $\alpha$ | 10 | Maturation rate of SFGFP protein | 1/sec |
| $k_i$ | 11 | Translational initiation rate of SFGFP | 1/sec |
| $r_{m2}$ | 12 | Maturation rate of $R_2$ | 1/sec |
| $r_{m1}$ | 13 | Maturation rate of $R_1$ | 1/sec |
| $p_c$ | 14 | Relative crosstalk level 1 experiences from level 3 | N/A |
| $P_t$ | 15 | Probability of that an attenuator auto terminates itself | N/A |
| $N_1$ | 16 | Number of tandem attenuator in level 1 | N/A |
| $N_2$ | 17 | Number of tandem attenuator in level 2 | N/A |

**Table 2.**
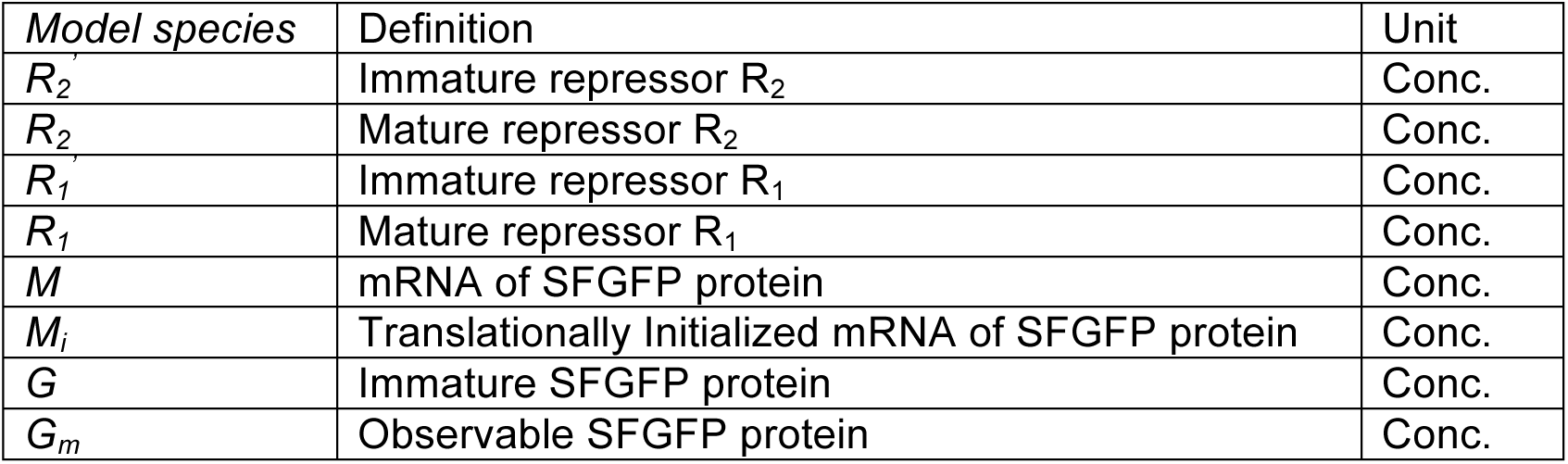
Model species.

### Parameter Estimation Through Sensitivity Analysis-Based Experimental Design

Our next goal was to determine parameters that would give quantitative agreement between our model and measurements of the final output fluorescence of the RNA transcriptional cascade. Parameter estimation is a highly non-trivial problem, as parameters are complex functions themselves of many detailed biochemical reactions. For example, transcription rate is commonly modeled as a single parameter with units of nucleotides/sec, when in reality it encompasses many separate processes including polymerase-promoter recognition, open-complex formation, escape probability and non-uniform elongation rates^37^. The most common method for determining parameters is to find them in the literature, and optionally consider parameter ranges centered on literature values^23^. While useful, this approach has limitations, including the fact that literature parameters are often not measured in an experimental/cellular/construct context relevant to the functioning of the model circuit, parameter measurement may not be consistent with the approximations made by the modeling framework, and it may be impossible to find certain parameters especially in cases when a synthetic regulator does not exist in nature^16^. An alternative approach is to perform a series of specific experiments designed to isolate the measurement of each parameter. While effective^38^, this approach is difficult to scale as the size of the genetic networks grow.

Potentially more powerful are methods that can take into account the structure of the genetic circuit to design a minimal set of experiments that can be used to rapidly and quantitatively determine all parameters in a model. Such methods have been developed in the context of systems biology, which focuses on understanding a biological system’s structure and dynamics. Because natural biological networks are usually massive and full of unknown species and parameters, systems biologists have developed numerous methods to locate the most important components of a genetic network and identify the most sensitive parameters^2^. Here we adapted a technique based on parameter sensitivity analysis and used it to design experiments that would provide enough information to identify all unknown parameters in our RNA circuitry model (Figure 1).

The basis of this technique is to use sensitivity analysis to guide the design of experiments that can use the full time trajectory information of a genetic circuit’s output to determine multiple parameters at the same time (Figure 1). For a particular experiment modeled by a system of ODEs, sensitivity analysis is based on the sensitivity matrix, which describes how the time varying molecular concentrations of the genetic circuit in the experiment change in response to a change in the parameters of the model (see Methods)^39^. Since our experiments measure SFGFP expression, we focused our analysis on how sensitive predicted trajectories of SFGFP expression were to changes in the thirteen unknown parameters. Parameters with large magnitudes in the sensitivity matrix (highly sensitive) can then be determined by fitting them to make the model match the experimental data. By proposing different experiments, this procedure can be iterated until a panel of experiments is designed that together can be used to estimate all parameters of the model (Figure 1). We note that this procedure is particularly amenable to being used with TX-TL reactions since circuit DNA template concentrations can be easily varied to rapidly design a set of parameterization experiments.

To perform this procedure, we used an initial set of parameter guesses taken from the literature or manual fitting to calculate the sensitivity matrix from a proposed experimental design. Our goal was to determine a reduced set of TX-TL experiments that could be used to find all thirteen parameters of our RNA transcriptional cascade model. In order to strike a balance between TX-TL energy resource usage and potential bleaching effects of the fluorescence measurement, we targeted experiments that could be performed at 29°C for 100 minutes with fluorescence collection every 5 minutes. We first performed the sensitivity analysis on an experiment consisting of just the bottom level (L1) of the cascade (Figure 3A). The calculated sensitivity matrix for the subset of equations that model L1 revealed that the SFGFP fluorescence trajectory in this experiment is most sensitive to the parameters β_m_ (promoter strength), k_E_ (translational elongation rate), d_m_ (mRNA degradation rate), α (SFGFP maturation rate), and k_I_ (translational initiation rate) (Figure 3A). This recapitulates the relationship that steady-state SFGFP concentration should be related to the product of the transcription and translation rates divided by the degradation rates of mRNA and SFGFP.

**Figure 3.**
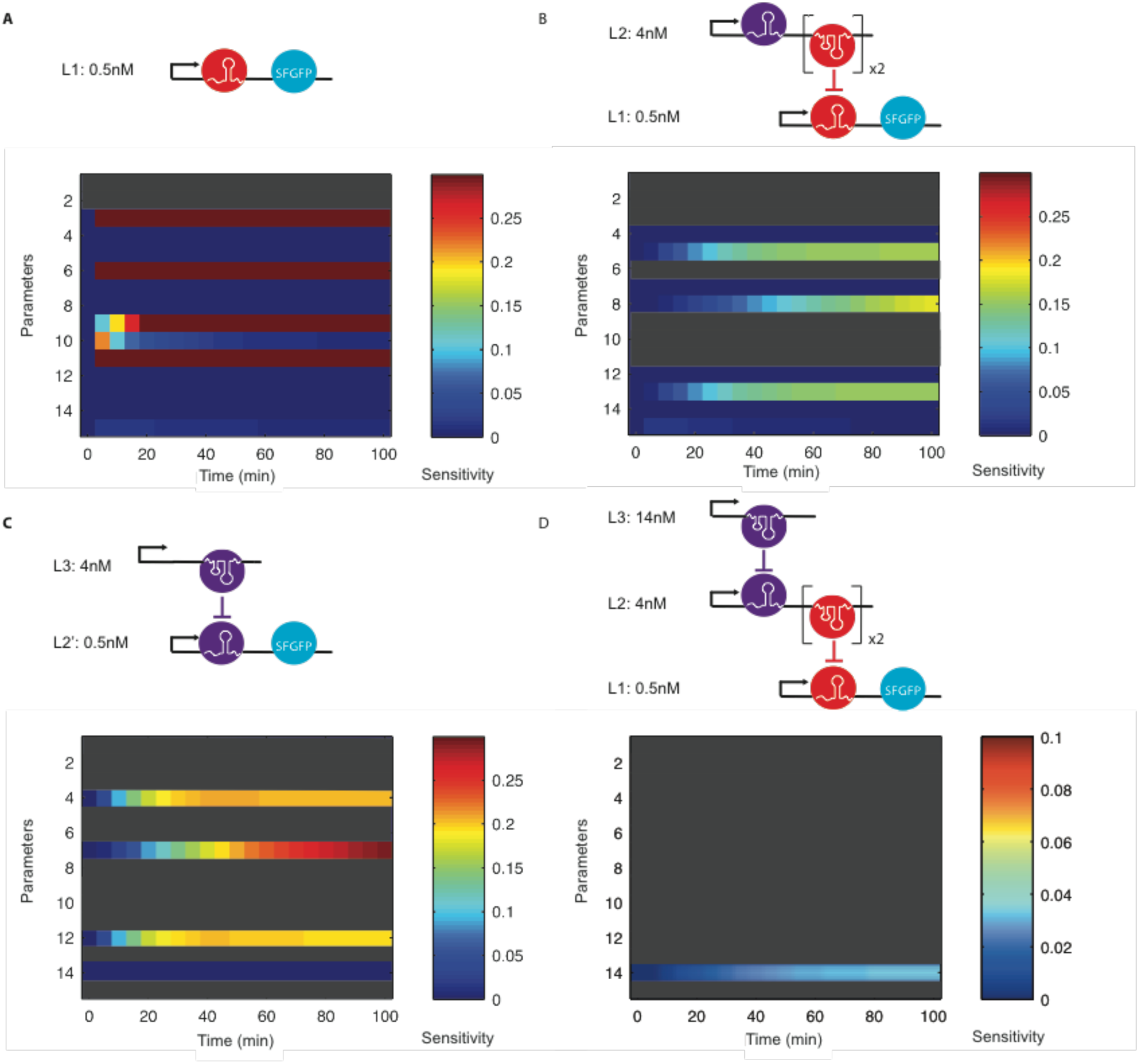
Sensitivity matrices for parameter identification experiments. Four parameterization experiments were designed based on sensitivity analysis to estimate all thirteen unknown parameters in our model (Figure 2). For each experiment, the constructs used are shown above the SFGFP portion of the calculated sensitivity matrix for that experiment. Parameters are numbered according to Table 1. Red/blue indicates high/low sensitivity, respectively. Note that time varying changes in parameter sensitivity indicates portions of the trajectories that are influenced by each parameter. Experiments were designed in order from (A) to (D), with parameters identified by previous experiments marked as grey rows. Since the same promoter was used on all constructs, β_1_ and β_2_, determined by multiplying β_m_ by an appropriate factor based on DNA template concentrations, are absent from the sensitivity analysis. (A) 0.5nM of L1 plasmid alone is able to identify five parameters: β_m_, k_E_, d_m_, α and k_I_.0.5nM of L1 and 4nM of L2 is able to identify four parameters: K_1_, d_1_, r_m1_ and P_t_. (C) 0.5nM of L2’ (Table S2) and 4nM of L3 is able to identify three parameters: K_2_, d_2_, r_m2_ (D) 0.5nM of L1, 4nM of L2, and 14nM of L3 is able to identify the last parameter, p_c_. The sensitivity color scale was changed in D to aid in visualization.

We next proposed an experiment that added a single layer of repression (Figure 3B). Since β_m_, k_E_, d_m_, α and k_I_ were already identified in the previous experiment, their rows in the SSM were all set to 0 so that the algorithm skipped identified parameters from its previous rounds and searched for the next most identifiable parameters. This second experiment was additionally able to identify 4 more parameters (*K*_*1*_, *d*_*1*_, *r*_*m1*_, *P*_*t*_) (Figure 3B). Successive iterations allowed us to find two additional experiments that allowed all thirteen parameters to be identified with a total of four TX-TL experiments (Figure 3C,D).

We next sought to use this designed set of experiments to estimate the parameters in our model. We first performed replicate TX-TL experiments with each of the plasmid combinations we designed, and collected the fluorescence time trajectories of the reactions. Using this experimental data, we then estimated parameters using an iterative fitting procedure that used each experiment in turn to find its designated parameters (see Methods). Rather than focus on a single set of optimal parameters, we considered variations of parameter values that can capture the natural variation in experimental conditions. To do this, we initially input sets of parameters, drawn from uniform distributions centered around an initial best guess, into the fitting procedure and optimized the values of each parameter within each input set (see Methods). This resulted in 10,000 sets of the thirteen estimated parameters, which allowed us to calculate mean predicted trajectories with 95% confidence intervals (Figure 4).

**Figure 4.**
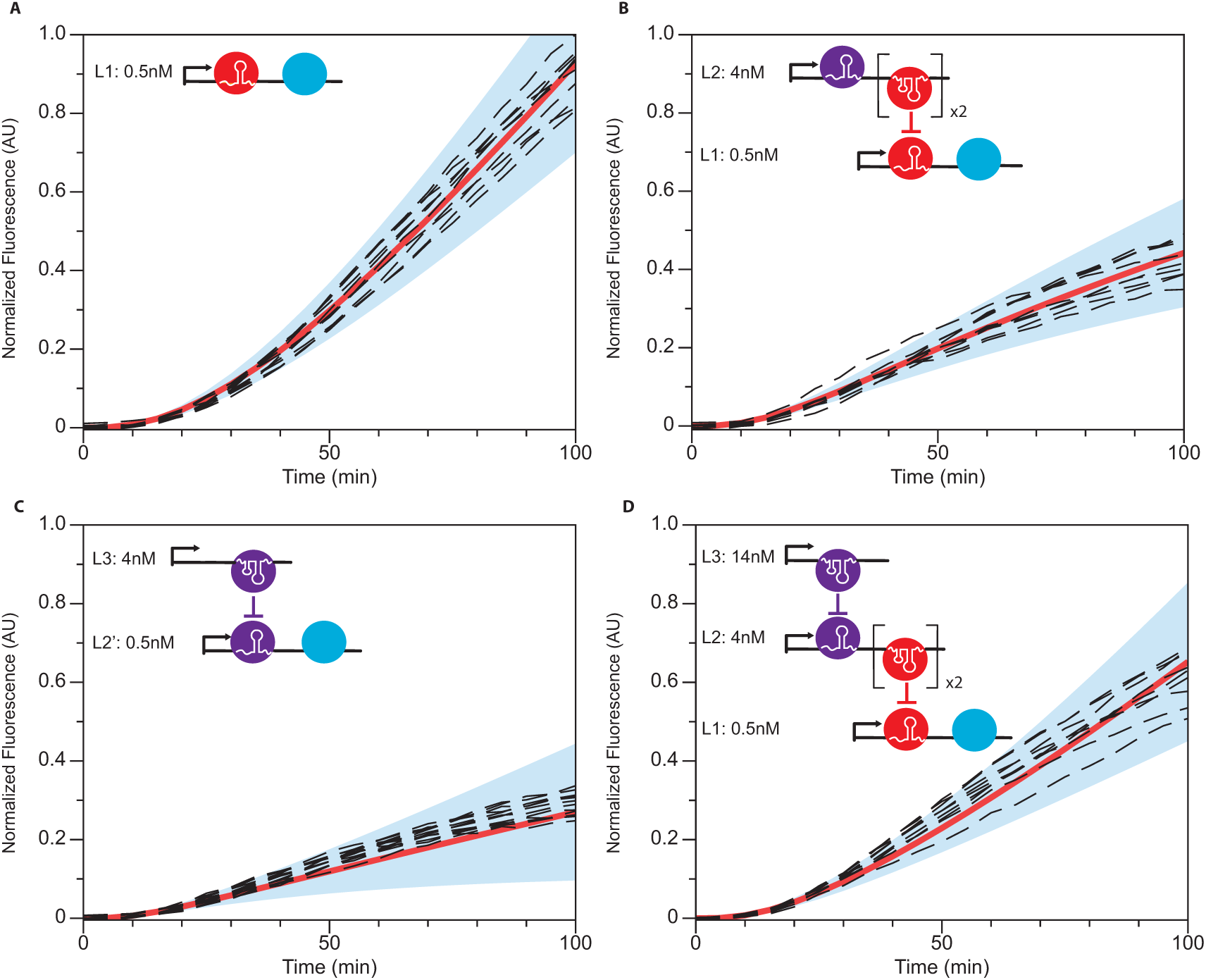
Validation of model simulations of parameter estimation experiments. Comparison of experimental trajectories of SFGFP fluorescence in TX-TL experiments (black dash lines) with simulated model predictions. Model simulated trajectories were generated by performing 1,000 simulations with parameters drawn from the set of 10,000 determined from the estimation procedure (see Methods). Experimental and model trajectories were normalized by the maximum observed experimental fluorescence (see Methods). The mean simulated trajectory (red line) is shown within 95% confidence intervals (blue region). The schematic of each experiment is shown in the upper left corner of each plot corresponding to the experiments in Figure 3.

As mentioned before, there were two R_1_ repressors encoded in L2 and one R_2_ repressor in L3 of the cascade, though they were both treated as single repressors in the model. Because the model was not given explicit information about the difference between these configurations, this gave us the opportunity to examine the estimated parameters to test the adaptability of our parameter estimation procedure to this type of model discovery (Table S4). Several estimated parameters showed that indeed this was the case. In particular, the repression coefficient of R_1_ (K_1_, mean 240.9) was significantly greater than the repression coefficient of R_2_ (K_2_, mean 132.0), indicating a weaker repression made by the double repressor configuration. This was actually consistent with previous *in vivo* characterization experiments, which showed that the L2 configuration containing an attenuator followed by tandem ribozyme-antisense coding regions was less efficient at repression than a single bare antisense^12^. In addition, the degradation rate of R_1_ (d_1_, mean 0.83 × 10^-3^ s^-1^) was also noticeably smaller than the degradation rate of R_2_ (d_2_, mean 2.08 × 10^-3^ s^-1^), showing that it is slower to degrade two repressors than it is to degrade one. Finally, the maturation rate of R_1_ (r_m1_, mean 3.21 × 10^-5^ s^-1^) was smaller than the maturation rate of R_2_ (r_m2_, 1.55 × 10^-4^ s^-1^), indicating additional processing steps are needed for the tandem repressor configuration. This makes intuitive sense given that ribozymes were included between the tandem R_1_ units, which must fold and cleave before R_1_ can properly function^12^.

To validate our method, we compared measured trajectories of SFGFP fluorescence for each experiment to simulated trajectories using 1,000 sets of randomly drawn parameters from the 10,000 parameter set distributions (Figure 4). This allowed us to compare the experiments to the mean simulated trajectory and 95% confidence intervals, which captured the range of trajectories predicted by our parameter distributions. As shown in Figure 4, each mean simulated trajectory accurately described the experimental observations, with almost every measured experimental trajectory lying within the simulation 95% confidence intervals.

These results validated our hypothesis that sRNA genetic circuits could be accurately modeled using simplified sets of ODEs with appropriate parameters. It also showed that our effective ODE model was able to quantitatively capture the complex biochemical conversions that take place within our model RNA circuit. Finally, it proved that our approach of estimating unknown parameters, using a minimal set of experiments designed from sensitivity analysis, was efficient, and produced accurate results. Conveniently these four TX-TL experiments could all be performed at the same time, allowing the simultaneous determination of all parameters in our model in a matter of hours. This approach is also highly generalizable to other ODE systems, and could be used to parameterize a broad array of synthetic genetic circuits.

### Model Predictions

Our ultimate goal of building a quantitative model was to accurately predict the behavior of a newly designed experiment. To do this, we designed six new experiments that varied the basic elements of the 3-level cascade, and compared model-simulated trajectories to experimental SFGFP fluorescence time course trajectories of these experiments (Figure 5).

**Figure 5.**
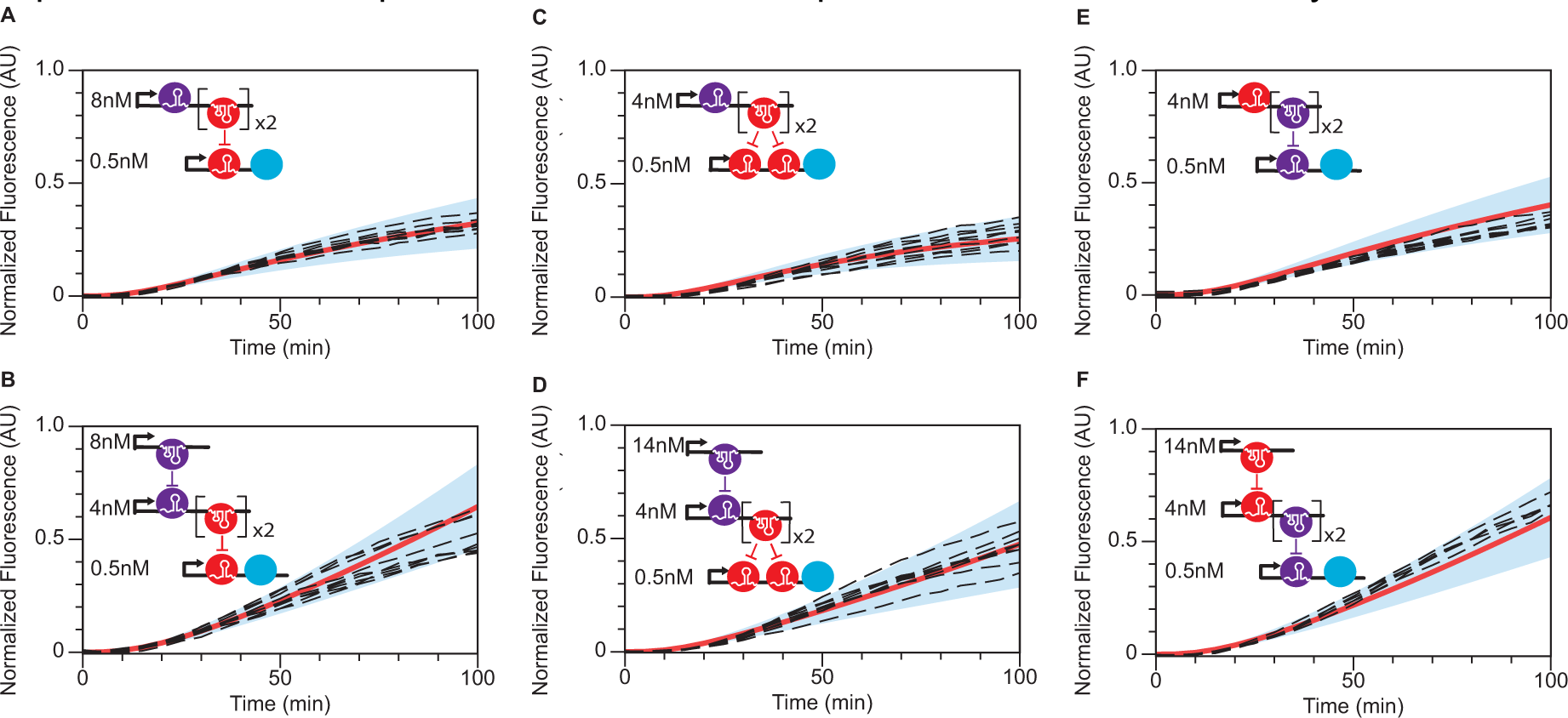
Model Predictions. Comparison of experimental trajectories of SFGFP fluorescence in TX-TL experiments (black dash lines) with simulated model predictions. Model simulated trajectories were generated by performing 1,000 simulations with parameters drawn from the set of 10,000 determined from the estimation procedure (see Methods). Experimental and model trajectories were normalized by the maximum observed experimental fluorescence (see Methods). The mean simulated trajectory (red line) is shown within 95% confidence intervals (blue region). The schematic of each experiment is shown in the upper left corner of each plot. Two level concentration prediction that varies the L2 plasmid concentration from Figure 3B. Three level concentration prediction that varies the L3 plasmid concentration from Figure 3D. (C) Two level tandem attenuator prediction. The experiment contains 0.5nM of a modified L1 plasmid expressing 2 tandem copies of A_1_ in front of SFGFP (L1T, Table S2), and 4nM of the L2 plasmid. (D) Three level tandem attenuator prediction containing the same constructs as in with an additional 14nM of L3 plasmid. (E) Two level swap prediction. The experiment contains 0.5 nM of a modified L1 plasmid that expresses A_2_ in front of SFGFP (PL1, Table S2), and 8nM of a new L2 plasmid expressing A_1_ followed by R_2_ (PL2, Table S2). (F) Three level swap prediction containing the same constructs as in (E) with an additional 14nM of a new L3 plasmid that expresses R_1_ (PL3, Table S2).

We first aimed to test the ability of the estimated parameters to capture changes in DNA template concentration of the cascade. Two experiments were designed that varied the concentration ratio of a single repressive connection of the cascade (Figure 5A), and the amount of antisense sRNA expressed from L3 in the full cascade (Figure 5B). Model predictions were made by running simulations using our estimated parameter sets with the parameters β_1_ and β_2_ multiplied by factors that took into account the change in DNA concentrations in the new experiments. In both new experiments, the average model predictions accurately matched the observed experimental trajectories, with all experimental trajectories falling within the 95% confidence intervals of the simulated trajectories (Figure 5A, B). In the single repression case of increasing L2 DNA, more repression was modeled and observed compared to the trajectories from the training experiment as expected (Figure 5A vs. Figure 4B). In the full cascade example, there were two sets of experimental trajectories observed, with the model showing predictions that captured the average behavior of these sets (Figure 5B).

Next we tested the ability to make predictions on RNA transcriptional cascades that included regulatory parts not involved in the training of any parameters. In particular, we designed an experiment to include a tandem attenuator in front of the L1 SFGFP reporter. Since the tandem attenuator is more sensitive to antisense RNA concentration, we expected both a single repressive connection and the full cascade to show an overall lower SFGFP signal throughout the trajectory^15^. This was indeed observed (Figure 5C vs. Figure 4B and Figure 5D vs. Figure 4D). Furthermore, we found that the model predictions for these new circuit variants were in strong agreement with the experiments, with the average predictions accurately capturing the observed trajectories, for which all but one fell within the 95% confidence intervals of the simulated trajectories (Figure 5C,D). It is important to note that the tandem attenuator regulatory part was not included at all in the parameterization experiments. Rather it was modeled by squaring the repressive function of the single-attenuator (Figures 2C, S1C, S2B). This shows how our model is easily extensible, and that accurate predictions can be made from a limited set of parameterization experiments when modular parts are used to construct synthetic circuits.

Finally we tested the ability of our model to predict rewiring of the cascade elements. We designed two experiments that swapped the order of the R_1_/A_1_ and R_2_/A_2_ repressive sRNA/target attenuator pairs in both a single repressive connection and the full cascade (Figure 5E,F). This was a challenging prediction since the genetic contexts of R_1_ and R_2_ changed in the swapped configuration between tandem ribozyme-antisense constructs and single antisense expression. As we observed in the parameterization experiments, this context change causes large changes in the repression parameters for these antisense sRNAs (Table S4). Therefore, we needed a way to easily estimate these new parameters using our previous information and as few as possible new parameterization experiments. To do this, we made several assumptions: (i) if the repression ratio of single copies of the two repressors, K_1s_ and K_2s_, is f (f= K_1s_/ K_2s_), then the repression ratio of tandem copies of the two repressors, K_1d_ and K_2d_, is also f (f= K_1d_/ K_2d_); (ii) the crosstalk strength of the swapped construct is the same as the crosstalk strength of the parameterization construct; and (iii) two different single repressors would have similar degradation and maturation rates, so that in the swap construct these two sets of maturation and degradations rates are unchanged. These assumptions allowed us to re-use almost all parameters except K_1s_ and K_2d_. Since K_2s_ and K_1d_ were already estimated in the parameterization procedure, we used one additional simple experiment to estimate K_1s_ (Figure S5). This allowed us to calculate f, which could then be used to calculate K_2d_ by our assumptions. Finally K_1s_ and K_2d_ were used to make the prediction of our new design with the rest of the previously determined parameters. Comparisons between the average simulated trajectories were again in strong agreement with the observed SFGFP fluorescence trajectories for these experiments, with all trajectories falling within the 95% confidence intervals of the experiments (Figure 5E,F).

Overall we found that our model and its estimated parameters are capable of performing quantitative predictions of new sRNA circuits. This demonstrated its potential for aiding in the design of circuits for synthetic biology. In addition, we showed that our model is extensible, and can incorporate new genetic parts with limited additional characterization experiments.

### Understanding TX-TL Batch Variation Using Model Parameterization

Given the simplicity and convenience of the four parameterization experiments, we wanted to investigate if these experiments could be used as a way to study batch-to-batch variation in TX-TL extract performance. In previous work, we observed differences between sRNA circuitry characterization time courses when different batches of TX-TL were used^15^. This was hypothesized to be due to different concentrations of molecular machinery in the extract that could impact the overall transcription, translation and degradation rates that influence circuit expression. Since our parameter estimation procedure establishes quantitative values for each of these key rates, we thought that comparing estimated parameters from two different batches of TX-TL would more precisely reveal the specific differences between the batches.

To test this idea, we performed the same parameter estimation procedure with a separate TX-TL batch. This yielded the same estimates of all thirteen parameters in our model, which were shown to accurately model the new experimental SFGFP circuit characterization trajectories (Figure S6). Although these two batches showed comparable quality, we noticed that batch A had slightly lower GFP fluorescence trajectories compared to batch B in three out of four parameterization experiments (Figure S7). To analyze this batch-to-batch difference, we compared the distributions of the thirteen parameters derived from these two sets of experiments (Figure 6, Table S4).

**Figure 6.**
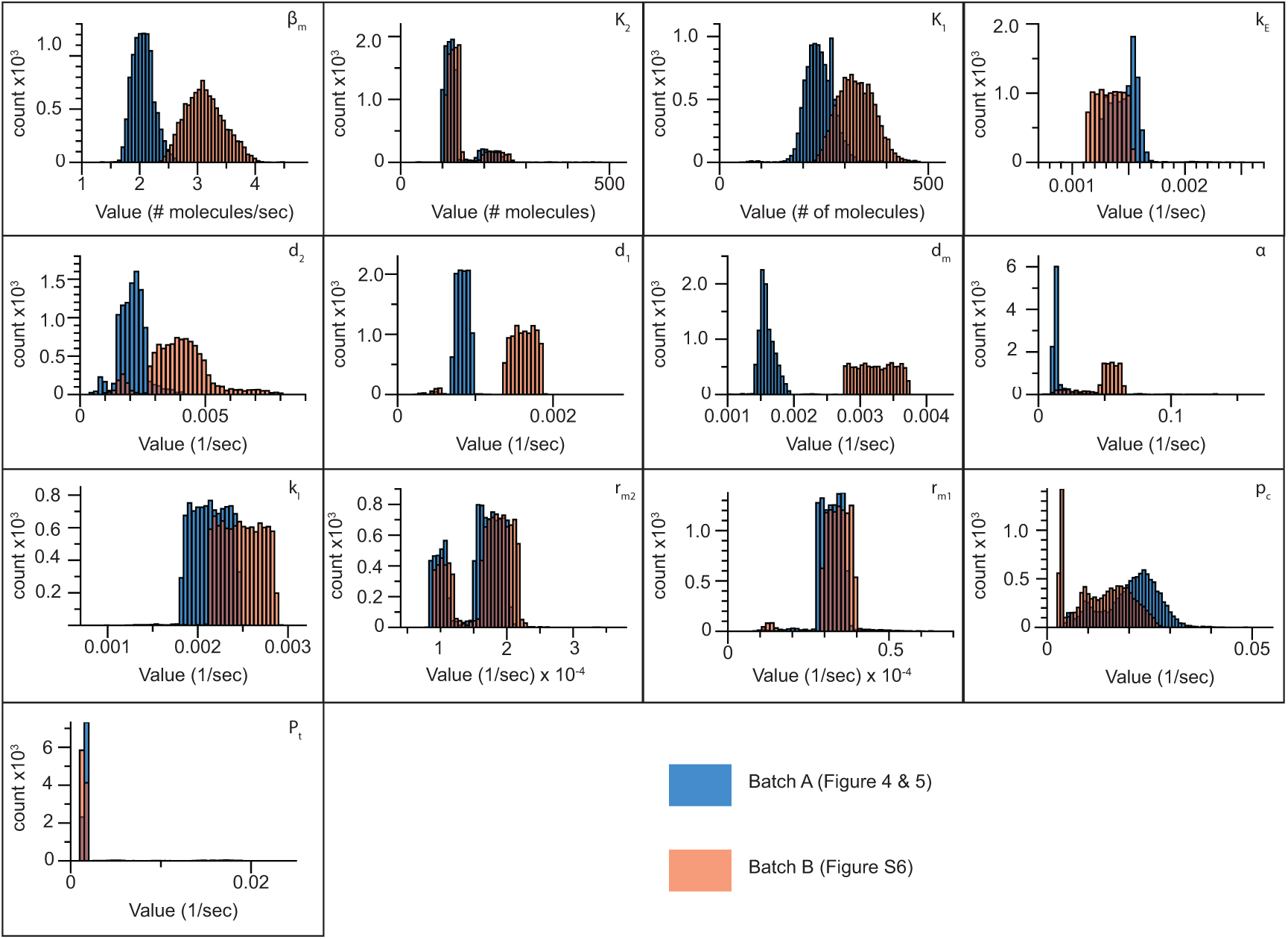
Parameter distributions from two independent TX-TL batches. Estimated parameters from batch A (blue) (Figure 4) and batch B (red) (Figure S6). Histograms are composed of 10,000 sets of parameters fit from the parameterization experiments performed in each batch.

Interestingly, we found that several parameters showed large differences between the two batches. These included the transcription rate β_m_, RNA degradation rates (d_2_, d_1_, d_m_) and SFGFP maturation rate α. Specifically, batch A had lower values for all of these parameters, suggesting that batch B had faster transcription, RNA degradation and GFP maturation. This could be explained by key differences in molecular machinery concentrations between these two batches that are known sources of variability in extract preparation^40^. Furthermore, these results demonstrated that our rapid parameter estimation procedure can be used to generate hypotheses about specific differences between TX-TL extract preparations, which could become increasingly important given the emerging new applications for these systems as molecular diagnostic platforms^41^.

## Conclusion

In conclusion, we have developed an effective quantitative model for sRNA transcriptional circuits and demonstrated its accuracy using a 3-level sRNA repression cascade as a test case. Our results showed that we were able to capture key features of this mechanistically sophisticated circuitry using only 8 ODEs with thirteen unknown parameters. To determine these parameters, we used sensitivity analysis to design four simple experiments that can be performed in parallel using cell-free TX-TL extracts to estimate all thirteen parameters. Finally we used our model along with estimated parameters to predict the time course dynamic trajectories for new network designs that used parts that were not included in parameterization experiments. We also showed that our model was easily extensible to include new parts with a limited number of additional characterization experiments. In all cases, our models were able to accurately reproduce the experimental results.

Interestingly, the process of constructing, parameterizing and validating our model of sRNA transcriptional circuitry revealed several new features of TX-TL systems and the sRNA regulators used. In particular, we found that there was a need to introduce a TX-TL pre-incubation step in order for computational analysis to match observed experimental trajectories. This is presumably due to certain processes that need to occur for the extract system to become active. In addition, we also discovered that the sRNAs used in our circuitry do not appear to be immediately active once synthesized, and we had to include equations that captured a maturation process to accurately model experimental SFGFP trajectories.

Several aspects of this work are significant. First, this represents the first validated computational model of synthetic RNA transcriptional genetic circuitry. Combined with other models that capture the effects of sRNA translational regulation^42^, this work helps lay the foundation for CAD tools that can incorporate RNA regulators into synthetic circuitry design^20^. We anticipate this to be more important as researchers increasingly turn to RNA-mediated gene regulatory systems for controlling gene expression in biological systems.

Second, the sensitivity analysis-based parameterization procedure is completely general, and could find wide use for establishing parameters for many synthetic regulators and circuits. In particular, the same exact procedure could be used to find parameters for the wide array of synthetic regulators now at our disposal^6^, including the exciting CRISPRi-based regulatory mechanisms. While we used the speed and experimental convenience of TX-TL reactions to perform our circuit parameterization experiments, this method should be extensible to models that capture the behavior of networks *in vivo*. In addition, this methodology could become even more powerful when coupled to different methods for implementing parameterization experiment design. For example, while we considered experimental design in terms of including different concentrations of DNA templates in the TX-TL reactions, one could easily imagine applying optogenetic control of component activity to design complex experiments that could allow even more efficient parameter estimation^38^. In these systems, different patterns of input light could be used to drive the system in specific ways designed to obtain the maximum number of parameters in the minimum amount of time.

Finally, we showed that our overall modeling and parameterization procedure offers a new approach for studying the underlying causes of batch-to-batch variation in TX-TL systems. In particular we found that different batches of TX-TL led to different distributions for key parameters that are directly related to the concentrations of core transcription/translation machinery in the batches (Figure 6). Understanding the basis of TX-TL variation will become increasingly important as these systems find wide use for a variety of applications including as metabolic production systems^43^, for rapid prototyping and characterization of genetic circuits^15,35,40^, and for new types of molecular diagnostics^41^.

## Methods

### Sensitivity Analysis to Design Parameterization Experiments

Our model for RNA circuitry consists of a set of ordinary differential equations that describe the time varying rate of change in the concentrations of the molecular species that participate in the circuitry, x_i_(t) (Figure 2). These equations are parameterized by a set of parameters, *p*_*j*_, that we want to estimate by fitting model predictions to a small set of experiments. These experiments were designed through an iterative process of sensitivity analysis on the set of model equations (Figure 2).

An individual experiment was considered to be a TX-TL reaction containing a subset of the DNA constructs encoding the full three-level sRNA transcriptional cascade at defined concentrations. Each such experiment produces a measureable trajectory of SFGFP fluorescence as a function of time, and can be modeled by the subset of equations that describe the gene expression processes from the included DNA. For example, if the TX-TL reaction contains only L1, then only the last four equations in Figure 2 need to be used with *R*_*1*_=*R*_*2*_=0. After a specific experiment (subset of DNA) was proposed, the next step was to assess which parameters were ‘identifiable’ from this experiment, which is closely linked with parametric sensitivity analysis. Here we used the procedure proposed by McAuley and coworkers^44^ to first calculate and analyze the sensitivity coefficient matrix for the proposed experiment as follows.

For each experiment, the sensitivity coefficient matrix z_ij_(t), is a time-varying matrix that encapsulates how sensitive the concentration of the molecular species x_i_ is to a change in the parameter *p*_*j*_

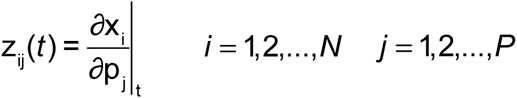

Here P denotes the number of parameters and N denotes the number of molecular species. If we write the model equations generally as

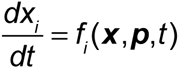

then it can be shown that Z_ij_(t) are the solutions to a set of differential equations given by

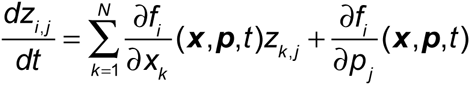

which are subject to the initial condition *zij* (0) = 0. Since our only observable in the TX-TL experiment is SFGFP, we focused specifically on *z*_*SFGFP,j*_*(t)* to determine which parameters were identifiable in the experiment.

To begin the experimental design process, we first determined a set of parameters that closely matched experiments by hand-fitting the parameters against SFGFP trajectories measured from TX-TL reactions containing subsets of the cascade DNA elements, using initial guesses based on the literature findings^34,45-47^. We next proposed the simplest experimental design (a fixed concentration of the bottom level L1) to the sensitivity analysis procedure. Using the hand-fit parameters, *z*_*SFGFP,j*_*(t*_*k*_) was calculated by solving the equations shown in Figure 2 using Matlab over a set of discrete time steps, *t*_*k*_, and then scaled by multiplying by *p*_*j*_*/x*_*SFGFP*_*(t*_*k*_). Identifiability was then performed according to McAuley^44^.This was done by finding the column of this matrix that had the biggest magnitude (indicating the most sensitive parameter), calculating a residual matrix which removed this column and controlled for correlations between parameters, and iterating this procedure on the resulting residual matrix until a threshold was reached on the largest remaining column magnitude. In this way a set of parameters was determined that maximally influenced the modeled trajectory of the proposed experiment (Figure 3A).

After performing this procedure on the simplest experiment (L1), we proposed a further experiment and performed the same analysis, except that parameters already identified by previous experiments were marked as ‘determined’ by setting their columns in the sensitivity matrix to 0. Rounds of experimental design and sensitivity analysis were performed until all 15 parameters were able to be identified by four TX-TL experiments (Figure 3).

### Parameter Estimation

Parameters were estimated from each designed experiment by fitting the identifiable parameters of that experiment to measured SFGFP expression time trajectories. The parameter estimation problem is given by:

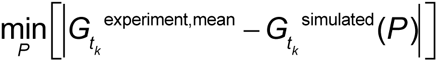

where 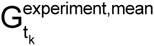 denotes the average value of the experimentally observed SFGFP expression at a certain time t_k_. The vector P contains all of the identifiable parameters in the experiment being analyzed. 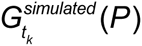 denotes the model simulated SFGFP expression at time *t*_*k*_. For a specific experiment, optimal P vectors were found using the *Matlab* function *fmincon*. Certain parameters were constrained to lie within specific values. For example, the repression concentrations K_1_ and K_2_ were constrained to be close in magnitude to each other based on the known similarity in repression of the two sRNA transcription repressor variants^12^. A complete list of constraints used is in Table S3.

Sets of identifiable parameters were estimated from the corresponding experimental trajectories in turn. We first used the estimation procedure to optimize the initial guesses used in the identifiability analysis above. To do this, we constructed uniform distributions (100 points) around each parameter value (± 15%), generating 100 sets of parameters. Each set of parameters served as a different starting point for finding optimal parameter estimates. Each estimate set was found by sequentially applying the fitting procedure above to each of the designed experiments, only fitting the identifiable parameters for that experiment. Fit parameters from one experiment were then used to update and replace the initial guesses before moving on to the next experiment until each of the 13 parameters was fit from the four experiments. This was repeated for each initial set of parameters to produce 100 sets of estimated parameters. We then chose the set that produced the closest simulated trajectory compared to the experimental data and used this as the guessed parameter set for the next iteration. We repeated this process 10 times. The final optimal parameter set was used in the same procedure to generate 10,000 sets of estimated parameters, which were then subject to the analysis outlined in the main text.

To plot our results, we scaled 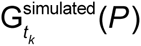 by the observed experimental values according to:

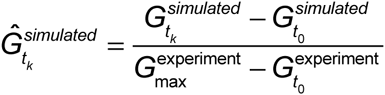

where 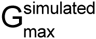 and 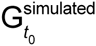 are the SFGFP expression level at any time *t*_*k*_ and initial time *t*_*0*_, respectively, and 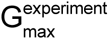 is the maximum experimental data point of the entire trajectories and 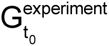 is the experimental data point at t=0. 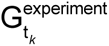 was also scaled as:

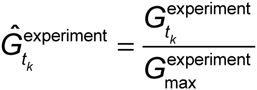

where 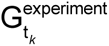 is the experimental expression of SFGFP at any time t_k_. In order to make all trajectories comparable, all experimental and modeled trajectories were scaled by the experimentally observed SFGFP fluorescence value of the first parameterization experiment at 100 minutes (Figure 4A).

### Plasmid construction and purification

A table of all the plasmids used in this study can be found in Supporting Table S2, with key sequences found in Supporting Table S1. The pT181 attenuator and repressor plasmids, pT181 mutant attenuator and antisense plasmids, and the no-antisense control plasmid were constructs pAPA1272, pAPA1256, pAPA1273, pAPA1257, and pAPA1260, respectively, from Lucks *et al.*^12^. The second level of the cascade (JBL069) was modified from construct pAPA1347 from Lucks *et al.*^12^. The double attenuator and modified level 2 constructs for prediction were created using Golden Gate assembly^48^. Plasmids were purified using a Qiagen QIAfilter Plasmid Midi Kit (Catalog number: 12243) followed by isopropanol precipitation and eluted with double distilled water.

### TX-TL extract and buffer preparation

#### Extract preparation

Cell extract and reaction buffer was prepared according to Shin and Noireaux^49^ and Sun *et al.*^35^. In brief, E. coli BL21 Rosetta cells were grown to an OD600 of 1.5, pelleted via centrifugation, and washed with a buffer at pH 7.7 containing Mg-glutamate, K-glutamate, Tris and DTT. Lysis was performed via bead-beating, followed by centrifugation to remove beads and cell debris. The resulting supernatant was incubated at 37°C for 80 minutes and then centrifuged, to remove endogenous nucleic acids. The supernatant was dialyzed against a buffer at pH 8.2, containing Mg-glutamate, K-glutamate, Tris and DTT. The extract was then centrifuged, and the supernatant flash-frozen in liquid nitrogen and stored at -80°C. The cell extract for Batch A had a protein concentration of 30 mg/mL, and its expression was optimized via the addition of 1 mM Mg and 40 mM K. For Batch B: 30 mg/mL protein,1 mM Mg, and 80 mM K.

#### Buffer preparation

The reaction buffer was prepared according to Sun *et al.*^35^, and consists of an energy solution (HEPES pH 8 700 mM, ATP 21 mM, GTP 21 mM, CTP 12.6 mM, UTP 12.6 mM, tRNA 2.8 mg/ml, CoA 3.64 mM, NAD 4.62 mM, cAMP 10.5 mM, Folinic Acid 0.95 mM, Spermidine 14 mM, and 3-PGA 420 mM) and amino acids (leucine, 5 mM, all other amino acids, 6 mM). Extract and buffer were aliquoted in separate tubes (volume appropriate for seven reactions) and stored at -80°C.

### TX-TL experiment

TX-TL buffer and extract tubes were thawed on ice for approximately 20 min. Separate reaction tubes were prepared with combinations of DNA representing a given circuit condition. Appropriate volumes of DNA, buffer, and extract were calculated using a custom spreadsheet developed by Sun *et al.*^35^. Buffer and extract were mixed together and incubated for another 20 min at 37°C. DNA for the specific experiment was then added into each tube according to the previously published protocol^35^. Ten μL of each TX-TL reaction mixture was transferred to a 384-well plate (Nunc 142761), covered with a plate seal (Nunc 232701), and placed on a Biotek SynergyH1m plate reader. We note that special care was needed when pipetting to avoid air bubbles, which can interfere with fluorescence measurements. Temperature was controlled at 29°C. SFGFP fluorescence was measured (485 nm excitation, 520 emission) every five min for 100 min. Each reaction was run with a minimum of triplicate repeats, and repeated three times independently (minimum of nine total replicates). A ten μL sample of each TX-TL buffer and extract mixture was run together with each independent reaction as blank. All data points were thenprocessed with blank values subtracted.

## Supporting Information

Supporting Tables 1-4, Supporting Figures 1-7, Supporting Appendix 1. Software for performing the identifiability and parameter estimation procedures in this work is available free of charge via the Internet at https://github.com/LucksLab/Hu_RNA_Circuit_Parameterization_ACSSynBio_2015.

## Abbreviations

Transcription-translation (TX-TL), super folder green fluorescent protein (SFGFP), attenuator (Att), antisense (AS).

## Author Contributions

CH performed the experiments. CH and JDV wrote the software. CH, JDV and JBL designed the experiments, analyzed the data and wrote the manuscript.

## Acknowledgement

We thank Melissa Takahashi and James Chappell for assistance in cloning and helpful conversations during this work.

